## Supplementary figures and images for "POMC neurons control fertility through differential signaling of MC4R in Kisspeptin neurons"

### Supplemental figure 1

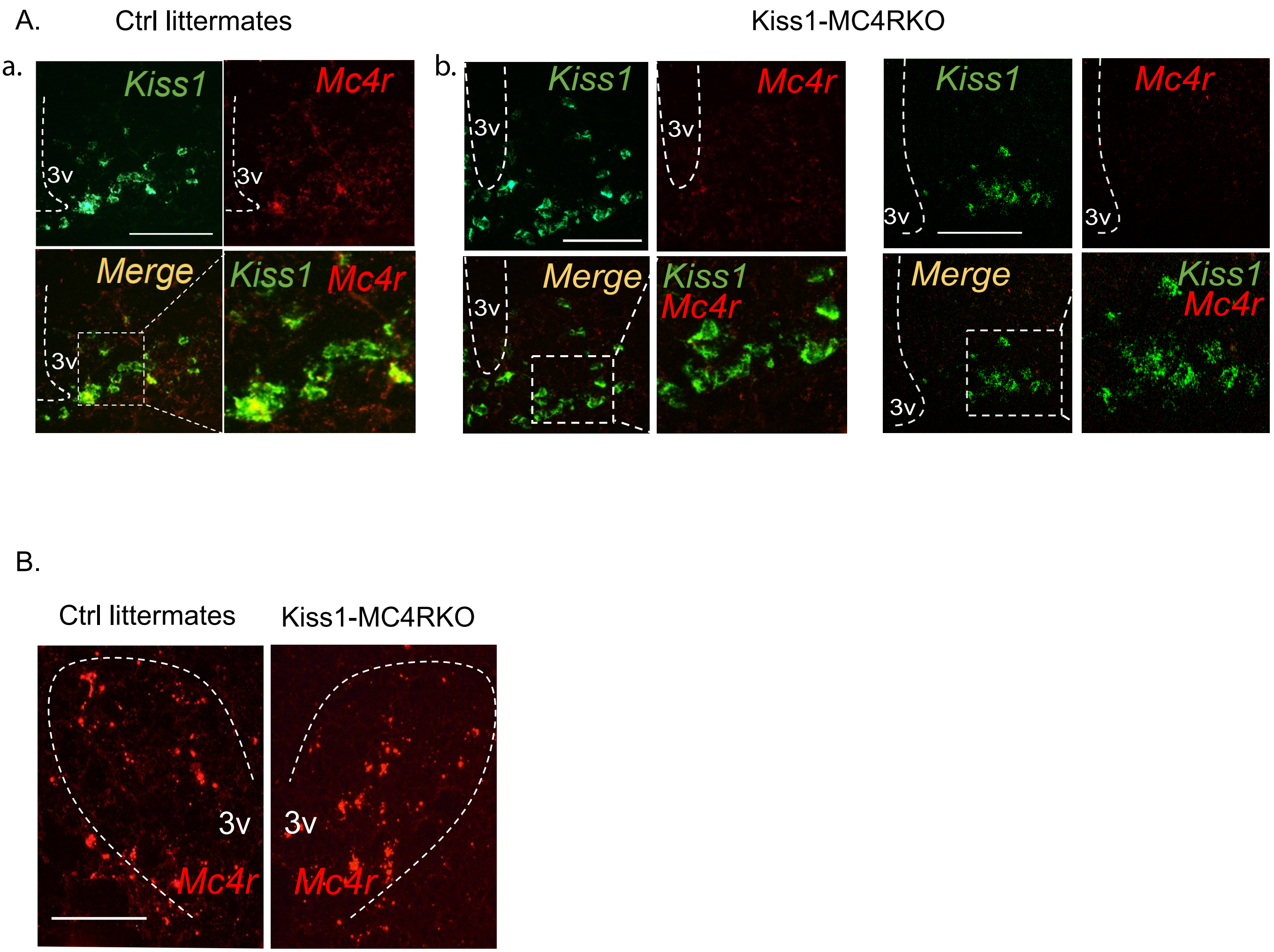

### Supplemental figure 2

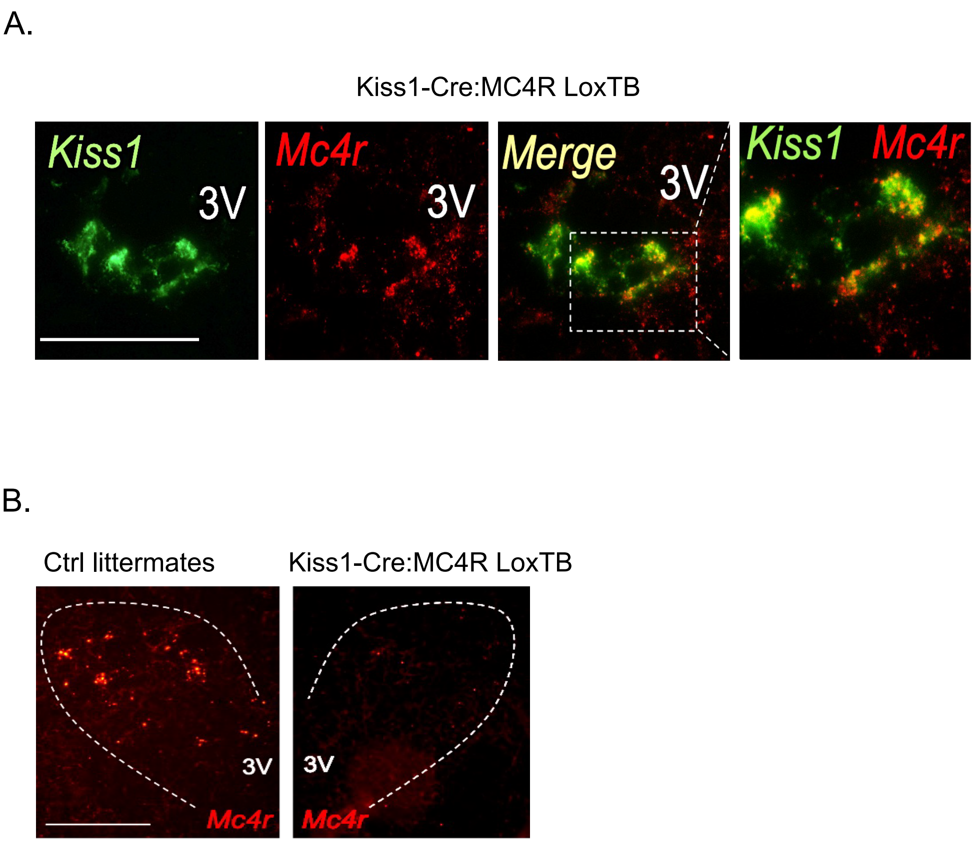

### Supplemental figure 3

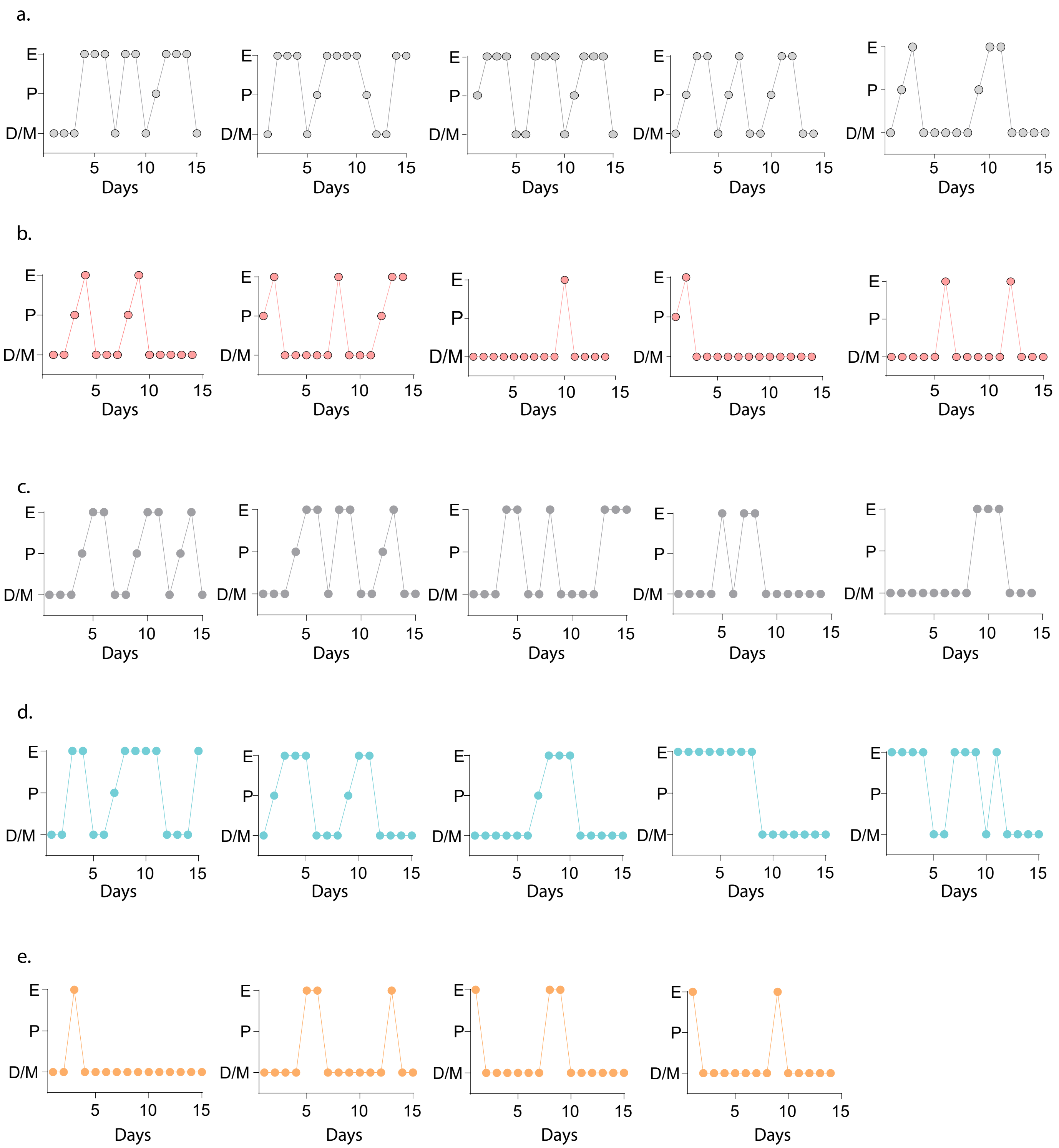
